# Comparative analysis of RNA interference and pattern-triggered immunity induced by dsRNA reveals different efficiencies in the antiviral response to *Potato virus X*

**DOI:** 10.1101/2024.02.07.579064

**Authors:** Khouloud Necira, Lorenzo Contreras, Efstratios Kamargiakis, Mohamed Selim Kamoun, Tomás Canto, Francisco Tenllado

## Abstract

Plant antiviral responses induced by double-stranded RNA (dsRNA) include RNA interference (RNAi) and pattern-triggered immunity (PTI), but their relative contributions to antiviral defense are not well understood. We aimed at testing the impact of exogenous applied dsRNA on both layers of defense against *Potato virus X* expressing GFP (PVX-GFP) in *Nicotiana benthamiana*. Co-inoculation of PVX-GFP with either virus-specific (RNAi) or nonspecific dsRNA (PTI) showed that nonspecific dsRNA reduced virus accumulation in both inoculated and systemic leaves. However, nonspecific dsRNA was a poor inducer of antiviral immunity compared to a dsRNA capable to trigger the RNAi response, and plants became susceptible to systemic infection. Studies with a PVX mutant unable to move cell-to-cell indicated that the interference with PVX-GFP triggered by nonspecific dsRNA operated at the single-cell level. Next, we performed RNAseq analysis to examine similarities and differences in the transcriptome triggered by dsRNA alone or in combination with homologous and heterologous viruses. Enrichment analysis showed an over-representation of plant-pathogen signaling pathways, such as calcium, ethylene and MAPK signaling, which are typical of antimicrobial PTI. Moreover, the transcriptomic response to the homologous combination had a greater impact on defense than the heterologous combination, highlighting quantitative differences between RNAi and PTI immune responses. In addition, we provide genetic evidence that *DICER-like2* and *4* as well as *Argonaute2* were positively involved in PTI-based defense against PVX-GFP, and that dsRNA-induced PTI was enhanced by salicylic acid signaling. Together, these results further our understanding of plant antiviral defense, particularly the contribution of nonspecific dsRNA-mediated PTI.

**IMPORTANCE:** Non-transgenic, RNA-based technologies based on topical application of dsRNA represent a promising approach for crop protection. Recent research has shown that in addition to the antiviral RNAi response, dsRNA activates also PTI defenses, contributing to plant immunity against virus diseases. However, little is known on the relative contribution of RNAi and PTI to antiviral defense. We found that while virus-specific dsRNA halted virus spread throughout the plant, nonspecific dsRNA reduced virus accumulation locally but was unable to prevent systemic infection in *Nicotiana benthamiana*. For the first time, a whole transcriptomic response to dsRNA in the context of a homologous and heterologous virus infection was examined, highlighting quantitative differences between RNAi and PTI immune responses. Our data suggest an unexpected connection between RNAi-related genes and PTI. We envisage that both sequence-specific RNAi and nonspecific PTI pathways may be triggered via topical application of dsRNA, contributing synergistically to plant protection against viruses.

## INTRODUCTION

Viruses are obligate intracellular pathogens that depend on a living cell to multiply and proliferate by hijacking the host cell metabolism and replication machinery. To protect themselves, plants have evolved a multi-layered immune system for defense against viruses, including innate immunity and RNA silencing, among others (1). RNA silencing or RNA interference (RNAi) is a regulatory mechanism evolutionarily conserved in most eukaryotes that relies on the sequence-specific degradation of targeted RNAs guided by complementary small RNAs (sRNAs) (2). In addition to its crucial activity in regulating plant development and growth, RNAi plays a main role in adaptive immunity against pathogens, including plant viruses (3). RNAi against viruses is triggered by double-stranded RNA (dsRNA) molecules that derive either from viral replication intermediates or from hairpin RNAs. The dsRNA is recognized by specific RNase type III-like enzymes designated in plants as DICER-like proteins (DCL), which cleave the dsRNA into sRNA duplexes of 21–24 nucleotides long (4). Then, the sRNAs bind to Argonaute (AGO) protein family members that constitute the core of the RNA-induced silencing complexes (RISC).

In *Nicotiana benthamiana*, several AGO and DCL proteins have been described with homology to those in *Arabidopsis thaliana* (5). DCL4 is the primary DCL component of antiviral defense against RNA viruses, as DCL4 mutants are more susceptible to different viruses (6). Specifically, it was shown that DCL4 played a moderate role in the inhibition of *Potato virus X* (PVX) in inoculated leaves, while DCL4 and DCL2 are both required to prevent systemic infection by PVX in Arabidopsis (7). DCL2 participates in the production of secondary sRNAs that reinforces systemic RNA silencing (8), and its role in antiviral defense is often masked by DCL4 activity (6). Several studies have suggested a broad involvement of AGO2 in defense against multiple RNA viruses (reviewed in 9). In particular, CRISPR-generated *N. benthamiana ago2* knockout plants were more susceptible to several viruses, including PVX (10).

Innate immunity largely relies on plant perception of the microorganism through the recognition of conserved pathogen-associated molecular patterns (PAMPs) by cellular pattern recognition receptors (PRRs), thus initiating the so-called pattern-triggered immunity (PTI) (11). Despite the broad knowledge on PTI against other pathogens, much less is known about PTI against plant viruses. Nonetheless, several findings suggest that PTI also plays an important role in plant–virus interplay. First, the exogenous application of dsRNA, a well-known PAMP in animal antiviral immunity (12), has been reported to be an elicitor of PTI against plant virus infection (13, 14). In contrast to RNAi, antiviral PTI triggered by dsRNA is independent of RNA sequence and may also be activated by non-viral sequences, e.g., green fluorescent protein (GFP)-derived dsRNA and the synthetic dsRNA analog polyinosinic-polycytidilic acid (poly (I:C)). However, plant PRRs involved in dsRNA recognition remain to be identified. Second, Arabidopsis mutants in the PRR coreceptor kinases *Somatic Embryogenesis Receptor-like Kinase (SERK)1*, *SERK3* and *SERK4* exhibit increased susceptibility to different RNA viruses (15, 16, 17). Third, plant viruses have acquired the ability to suppress PTI mechanisms via the action of viral effectors (18, 19). In particular, tobamovirus movement proteins suppressed the dsRNA-induced defense response leading to callose deposition at plasmodesmata (PD), thus enabling virus movement (17).

The antiviral PTI-like responses induced by dsRNA are somehow similar to those of antimicrobial PTI, and include the induction of ethylene (ET) production, the activation of mitogen-activated protein kinases (MAPKs), and the triggering of defense gene expression (11, 13, 14, 16). However, the perception of poly (I:C) did not lead to the production of reactive oxygen species (ROS), indicating differences between dsRNA-induced and microbial-induced PTI signaling pathways (13, 17). Antimicrobial PTI is regulated by phytohormones, with salicylic acid (SA), jasmonic acid (JA) and ET acting as central players in triggering the immune signaling network (20). For instance, it has been reported that SA and JA signaling were required for activation of PTI against *Pseudomonas syringae* induced by exogenous application of bacterial RNA, specifically rRNA, in Arabidopsis (21). Nevertheless, the contribution of phytohormone signaling to dsRNA-mediated PTI against infecting viruses remains largely uncharacterized.

Plant viruses represent a serious threat to agricultural production and food supply (22). Conventional breeding for introgression of major resistance genes even with new genetic marker technologies is a time-consuming and laborious process (23). Moreover, the heavy use of chemicals to control vectors transmitting plant viruses has a significant negative impact on human health and environment. Thus, alternative biotechnological approaches are needed to confer protection against virus diseases, which are more sustainable, environmentally friendly and positively perceived by society. Since the first example describing that exogenous application of dsRNA induced RNAi-based protection against RNA viruses (24), many studies have been documented reporting the topical application of dsRNA to confer protection against pests and pathogens in numerous plant species (reviewed in 25, 26). Given that dsRNA is the trigger of antiviral RNA silencing and can also act as a potent PTI elicitor, we envisage that both sequence-specific RNAi and nonspecific PTI pathways may be triggered via topical application of dsRNA, contributing synergistically to plant protection against viruses.

Plant antiviral responses include RNAi and PTI pathways, but their relative contribution to antiviral defense are not thoroughly understood. In this study, we have compared the impact of exogenously applied dsRNA on both these layers of defense, and explore its effect on the local and systemic accumulation of PVX expressing GFP (PVX-GFP) in *N. benthamiana*. Previous findings on PTI-responsive genes elicited by dsRNA were based on a limited number of candidate genes altered by microbial elicitors, which were selected from publicly available gene expression data (13, 14). To further investigate the dsRNA-triggered antiviral defense at genome*-*wide level, we performed next generation sequencing to examine similarities and differences in the alteration of the whole transcriptome triggered by dsRNA alone or in combination with homologous and heterologous viruses. After examining the RNAseq data, the contribution of phytohormones to dsRNA-mediated antiviral PTI was also studied. RNAi and PTI are often seen as two separate, non-connected mechanisms of antiviral defense. In Arabidopsis, several PTI hallmarks, i.e., MAPK activation and ET production, have been reported in dsRNA-treated DCL mutants, suggesting that PTI and RNAi act independently in antiviral defense (13). However, interference with virus accumulation triggered by nonspecific dsRNA in these plants has not been addressed. Thus, we have also investigated whether components of the RNAi pathway up-regulated by dsRNA were involved in PTI-based defense against PVX-GFP.

## RESULTS

### Differential reduction in virus abundance in virus-specific and nonspecific dsRNA-treated plants

*N. benthamiana* plants were inoculated with mixtures of PVX-GFP and bacterially produced dsRNA molecules homologous to either GFP (dsGFP) or the *Potato virus Y* (PVY) coat protein (CP) gene (dsPVY). The combination of PVX-GFP plus nucleic acid extracted from bacterial cells not expressing the dsPVY was used as negative control. A total of 12 plants from each treatment were tested in each of two independent experiments, and the subsequent viral infection was monitored by detecting GFP foci in the inoculated leaves at 7 days post-inoculation (dpi). There were fewer fluorescent spots or infection foci in the inoculated leaves treated with dsGFP than with the negative control, implying that virus-specific dsRNA partially interfered with PVX-GFP infection (Fig. 1A,B,C). However, there was no significant difference in the number of GFP foci between nonspecific dsRNA (dsPVY)- and control-treated leaves. Treatments with either dsGFP or dsPVY did not cause a significant change in the size of GFP foci (Fig. 1D). To examine the effect of dsRNA treatments on the accumulation level of PVX-GFP, we measured the relative amount of virus in inoculated leaves from dsGFP-, dsPVY- and control-treated plants at 7 dpi. Comparative analysis by quantitative RT-PCR (qRT-PCR) revealed that the level of PVX-GFP RNA in dsGFP- and dsPVY-treated leaves was reduced 5.4- and 2.2-fold respectively, compared to the control (Fig. 1E).

**Figure 1:**
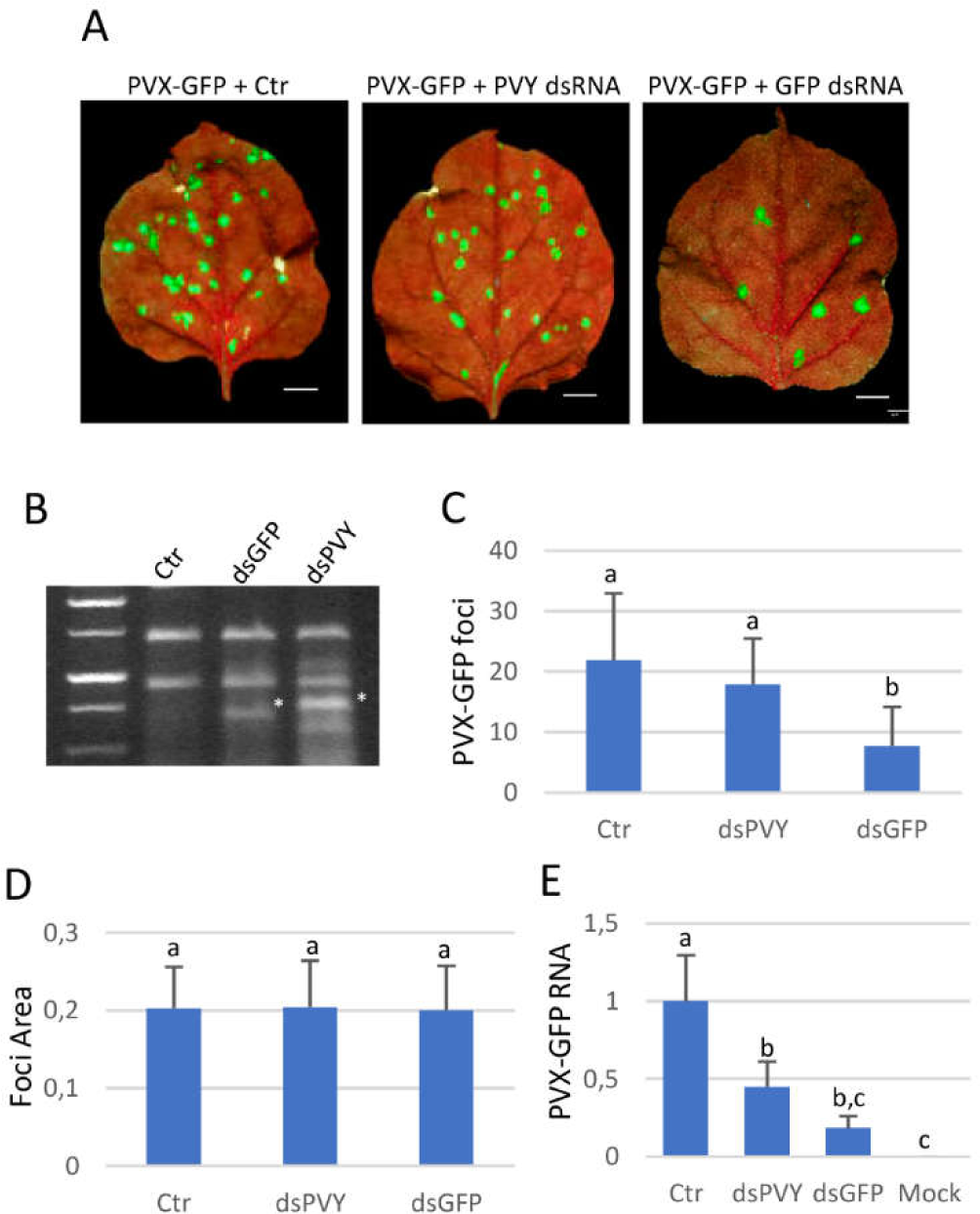
Virus-specific and nonspecific dsRNAs interfere differently with PVX-GFP local infection. *Nicotiana benthamiana* plants were inoculated with mixtures of PVX-GFP combined with nucleic acid extracts prepared from *E. coli* accumulating the dsPVY or dsGFP. (A) Representative inoculated leaves were examined under UV light at 7 days post-inoculation (dpi). Scale bar denotes 1 cm. (B) *E. coli* cultures transformed with L4440 encoding either a dsRNA consisting of 902 bp of the PVY CP gene (dsPVY) or GFP (dsGFP) were induced with IPTG and processed for total nucleic acid. The total nucleic acid extracted from bacterial cells not expressing dsRNA was used as a control (Crt). Samples were resolved by electrophoresis on 1% agarose gel. (C) Mean numbers ± standard deviation of infection foci on inoculated leaves of 12 plants (two leaves per plant) with the different treatments at 7 dpi. (D) Mean sizes ± SD of infection foci on inoculated leaves of 6 plants with the different treatments. (E) qRT-PCR was used to analyze the accumulation of PVX-GFP genomic RNA levels in the inoculated leaves at 7 dpi. Mock, mock-inoculated plants. Expression of the 18S rRNA gene served as a control. Data represent the means ± SD of 3 replicates, each consisting of a pool of 12 plants that received the same treatment. Different letters indicate significant differences determined by employing Scheffé’s multiple range test (*p* < 0.05). Experiments were repeated once more with similar results.

In independent experiments, the effectiveness of virus-specific dsRNA (dsGFP) or nonspecific dsRNA (dsPVY) in providing protection against PVX-GFP was tested by systemic infection assays. All of the 12 *N. benthamiana* plants inoculated with PVX-GFP plus control extract showed typical symptoms of mild mottling and GFP fluorescence in systemic leaves at 7 dpi. At the same time point, only 3 out of 12 and 10 out of 12 of plants treated with PVX-GFP plus either dsGFP or dsPVY showed GFP fluorescence, respectively (Fig. 2A). Northern blot analysis from RNA extracted from a pool of the upper leaf tissue from each treatment at 7 dpi revealed that accumulation of PVX-GFP genomic RNA was slightly lower in plants treated with PVX-GFP plus dsPVY than in control plants (Fig. 2B), whereas PVX-GFP accumulated below detection limits in dsGFP-treated plants displaying no symptoms of infection. This was corroborated by qRT-PCR analysis, where a 98 and 25% reduction in PVX-GFP RNA levels was quantified in plants treated with the virus-specific and nonspecific dsRNAs, respectively (Fig. 2C). Taken together, our findings show that nonspecific dsRNA induces a reduced inhibition of PVX-GFP accumulation in local and systemic leaves compared to dsRNA homologous to the targeted virus, suggesting that PTI responses are not strong enough to restrict systemic PVX-GFP infection.

**Figure 2:**
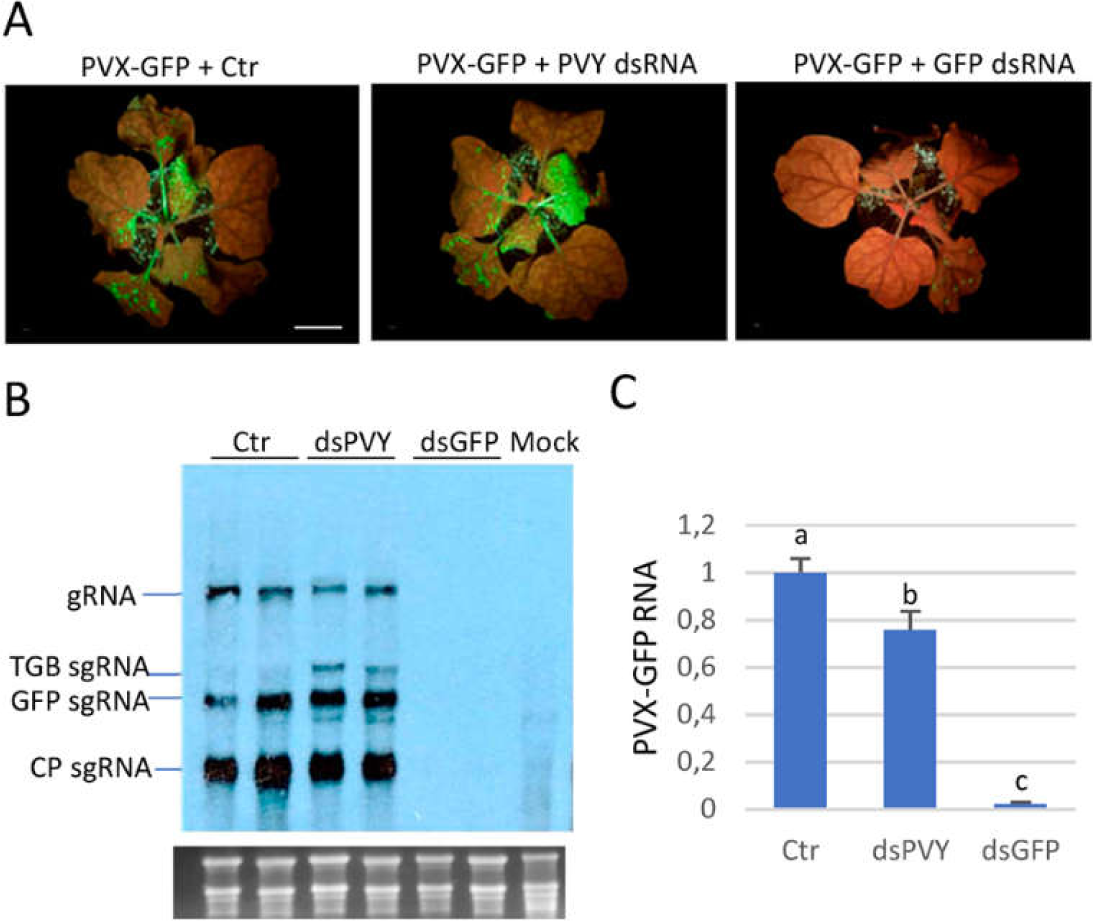
Systemic response of *Nicotiana benthamiana* plants to inoculation with mixtures of PVX-GFP combined with dsPVY, dsGFP or control (Ctr) extract. (A) Representative plants were photographed under UV light at 7 days post-inoculation (dpi). (B) Northern blot analysis of total RNA extracted from upper leaf tissues at 7 dpi. Two independent pooled samples were analyzed for each combination. Total RNA (5 µg) was hybridized with a probe complementary to PVX CP. PVX genomic (g)RNA and the major subgenomic (sg)RNAs, triple-gene-block (TGB) sgRNA, GFP sgRNA and CP sgRNA, are indicated. Ethidium bromide staining of rRNA is shown as loading control. (C) qRT-PCR was used to analyze the accumulation of PVX-GFP genomic RNA levels in the systemic leaves at 7 dpi. Expression of the 18S rRNA gene served as a control. Data represent the means ± SD of 3 replicates, each consisting of a pool of 12 plants that received the same treatment. Different letters indicate significant differences determined by employing Scheffé’s multiple range test (*p* < 0.05). Experiments were repeated once more with similar results.

### Nonspecific dsRNA causes inhibition of PVX-GFP replication

We next investigated the particular step of the viral replication cycle that was affected by nonspecific dsRNA. *N. benthamiana* plants were inoculated with mixtures of PVX-GFP plus either dsPVY, control extract or a bacterial PTI elicitor derived from flagellin (flg22) as a control, and samples were taken at an early time point after inoculation. At 4 dpi, there were no significant differences in the number and the size of GFP foci between the leaves treated with dsPVY or flg22 compared to control-treated leaves (Fig. 3A, and data not shown). Both PVX-GFP RNA and CP accumulated less in the inoculated leaves of dsPVY and flg22-treated plants than in the controls, as assayed by qRT-PCR and western blot analyses at 4 dpi (Fig. 3B,C).

**Figure 3:**
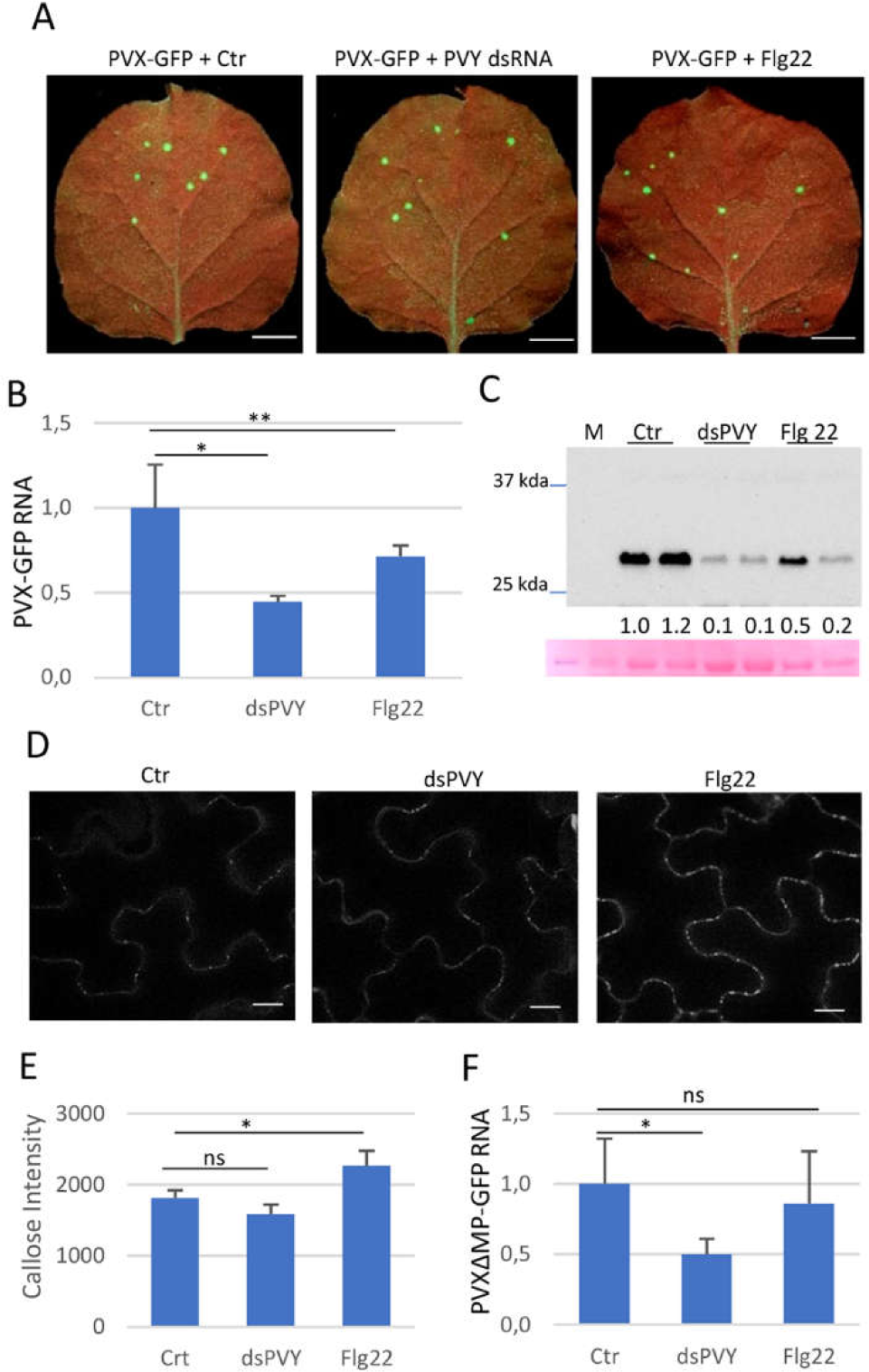
Nonspecific dsRNA elicits inhibition of PVX-GFP replication. Plants were inoculated with mixtures of PVX-GFP and either flagellin (flg22), dsPVY or control (Ctr) extracts. (A) Leaves were photographed under UV light at 4 days post-inoculation (dpi). (B) qRT-PCR was used to analyze the accumulation of PVX-GFP genomic RNA levels in the inoculated leaves at 4 dpi. Expression of the 18S rRNA gene served as a control. Data represent the means ± SD of 3 replicates, each consisting of a pool of 9 plants that received the same treatment. (C) Western blot analysis of plant extracts derived from inoculated leaves at 4 dpi, using antibodies against PVX CP. The panel below the blot is the membrane stained with ponceau S as control of loading. The intensity of each PVX CP band was quantified by densitometry analyses. (D) Callose spots at plasmodesmata (PD) were visualized upon aniline blue staining of epidermal cells in response to bacterial flg22, dsPVY or Ctr extracts. Photographs were taken 30 minutes after treatment with 1 µM flg22, 30 ng/ul of dsPVY or Ctr extracts. Scale bar, 10 μm. (E) Relative PD callose content in flg22-, dsPVY- and control-treated leaves. The mean values of callose intensities in individual PD (>200) measured in three leaf discs taken from two independent biological replicates per each treatment. (F) Accumulation of a frameshift mutant in the P25 movement protein of PVX-GFP (PVXΔMP-GFP), as assayed by qRT-PCR at 4 dpi. Asterisks indicate significant differences between treatments (Student’s t-test, *, *p* < 0.05; **, *p* < 0.1); ns: not significant. Experiments were repeated once more with similar results.

It has been reported that nonspecific dsRNA-induced immunity against *Tobacco mosaic virus* (TMV) restricts the progression of virus movement by triggering callose deposition at PD (17). To investigate whether PD callose deposition is involved in inhibition of PVX-GFP accumulation mediated by dsPVY in inoculated leaves, we used *in v*i*vo* aniline blue staining to quantify PD-associated callose in dsPVY- and flg22-treated leaves. Flg22 treatment caused an increase in PD callose intensity compared to leaves treated with control extract (Fig. 3D,E). By contrast, in our experimental conditions PD-associated callose in dsPVY-treated leaves remained at control levels, indicating that callose accumulation was not responsible of the inhibition of PVX-GFP accumulation caused by nonspecific dsRNA.

To test whether virus replication is affected by nonspecific dsRNA, we measured the accumulation of PVX-GFP RNA in leaves inoculated with a frameshift mutant in the P25 movement protein (PVXΔMP-GFP) which renders the virus unable to move between cells (27). As shown in Figure 3F, treatment of the leaves with dsPVY elicited a significant reduction in PVXΔMP-GFP accumulation compared to control-treated leaves at 4 dpi. Treatment with flg22 had not a significant impact on viral accumulation in PVXΔMP-GFP-inoculated leaves. Altogether, the unaltered size and number of infection sites together with the lack of callose accumulation in dsPVY-treated leaves suggested that the nonspecific dsRNA-triggered immunity against PVX-GFP was not linked to the reduced cell-to-cell movement of the virus, but to inhibition of virus replication.

### dsRNA causes transcriptome reprogramming in *N. benthamiana*

To identify early biological processes and pathways associated with dsRNA-based immunity, we conducted a global transcriptome RNAseq assay with RNA extracted at 4 dpi from *N. benthamiana* leaves inoculated with four different treatments, i.e., dsGFP alone (dsG), PVX-GFP combined with dsGFP (homologous combination, dsG_VG), wild-type (Wt) PVX combined with dsGFP (heterologous combination, dsG_V) and bacterial nucleic acid extracts not expressing dsRNA as a control (Ctr). Three biological replicates were analyzed for each treatment. Sequencing of 12 transcriptome libraries generated over 968 million mapped reads. On average 88.76% of the clean reads had quality scores at the Q30 level. The sequencing data are summarized in Table S1.

Principal component analysis (PCA) was used to sort RNAseq-based transcriptomic data according to gene expression levels. Two principal components explained 61.5% of the overall variance of gene expression profiles (35.8 and 25.7% for principal component 1 (PC1) and principal component 2 (PC2), respectively; Fig. 4A). PC1 seems to highlight a shift in gene expression between virus-infected samples, dsG_V and dsG_VG, and non-infected ones, dsG and Ctr, which suggests a differential transcriptional status in virus-infected leaves. In addition, PCA score plot revealed a clear separation between Ctr and samples treated with dsGFP, i.e., dsG, dsG_V and dsG_VG, indicating that treatment with dsRNA was the main source of variance underlying PC2.

**Figure 4:**
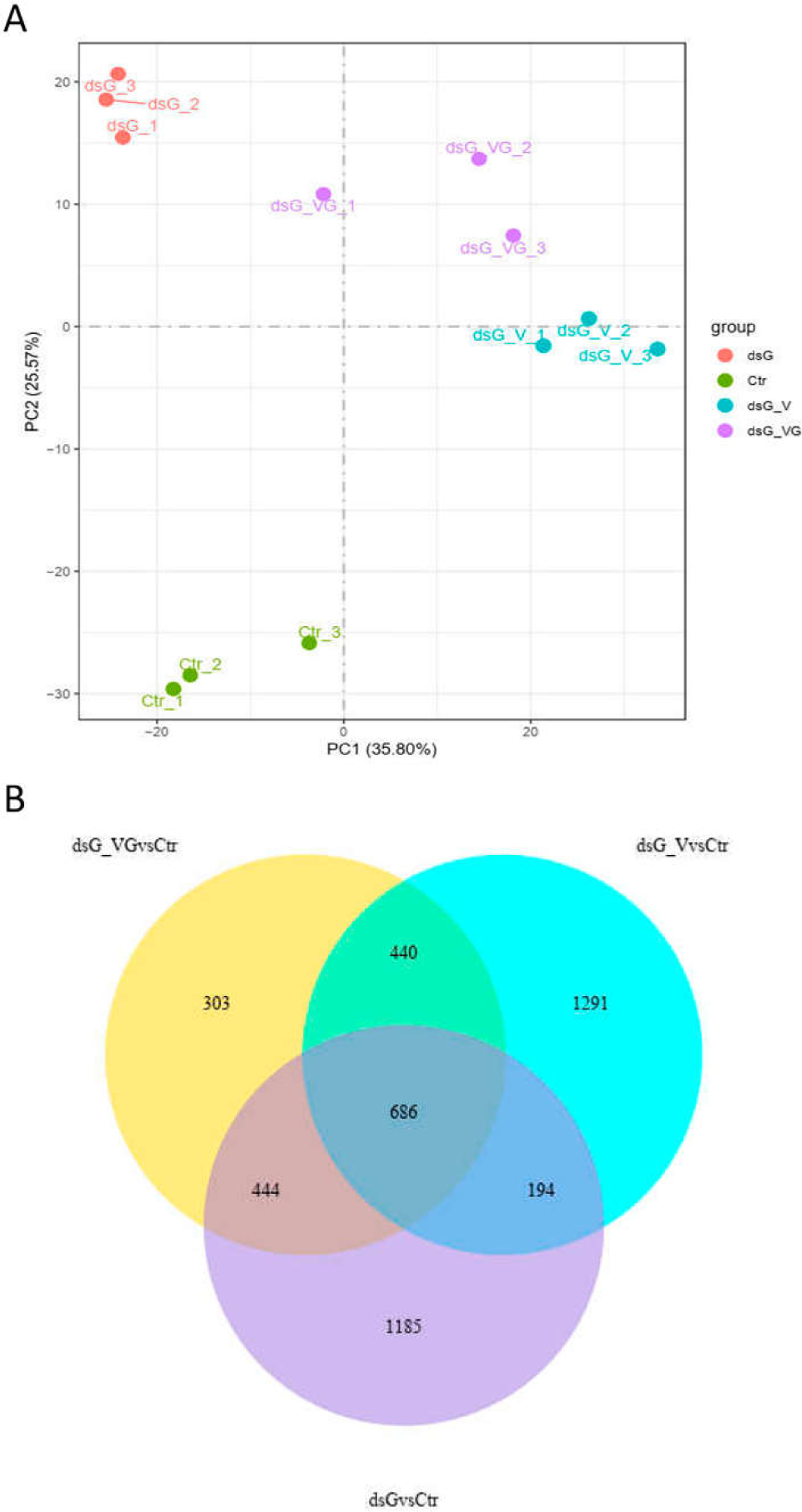
Transcriptional reprogramming associated to dsRNA and either homologous or heterologous virus infection. (A) Principal component analysis (PCA) of RNAseq data. The PCA was performed using normalized RNAseq data of differentially expressed genes (DEGs) between treatments at 4 days post-inoculation. Each biological replicate is represented in the score plot. The variance explained by each component (%) is given in parentheses. Treatments were as follows: dsGFP alone (dsG), PVX-GFP combined with dsGFP (dsG_VG), Wild-type PVX combined with dsGFP (dsG_V) and bacterial nucleic acid extracts not expressing dsRNA as a control (Ctr). PCA was performed using R package. (B) Venn diagrams displaying the number of DEGs with a |log2(Fold-change)| ≥ 1 and adjusted *p*-value ≤ 0.05 in the dsG vs. Ctr, dsG_VG vs. Ctr and dsG_V vs. Ctr pairwise comparisons.

Differentially expressed genes (DEGs) from paired comparisons are presented in Table S1 (adjusted *p*-value ≤ 0.05; |log2 ratio|≥1). We focused on comparisons between the expression profiles resulting from response to either dsRNA alone (dsG vs. Ctr), the homologous interaction between PVX-GFP and dsGFP (dsG_VG vs. Ctr), or the heterologous interaction between PVX and dsGFP (dsG_V vs. Ctr), to highlight quantitative and qualitative differences attributable to responses to dsRNA itself, RNAi and PTI, respectively. In the dsG vs. Ctr comparison, 2,509 DEGs, including 1,441 upregulated and 1,068 downregulated genes, were identified, whereas 2,611 (1,962 upregulated and 649 downregulated) and 1,873 (1,514 upregulated and 359 downregulated) DEGs showed significant changes in the dsG_V vs. Ctr and dsG_VG vs. Ctr comparisons, respectively (Fig. 4B).

The Kyoto Encyclopedia of Genes and Genomes (KEGG) pathway analysis was used to infer pathways significantly associated with DEGs in each dataset (Table S2). KEGG enrichment analysis (*p* < 0.05) allowed the identification of several terms, including Photosynthesis, Plant-pathogen interaction, Porphyrin and chlorophyll metabolism, Cutin, suberine and wax biosynthesis, Pentose and glucuronate interconversions, and MAPK signaling pathways, among others, overrepresented in the dsG vs. Ctr comparison (Fig. 5A). It is noteworthy that, among others, the KEGG terms Plant-pathogen interaction and MAPK signaling pathway were also overrepresented in the set of DEGs from dsG_V vs. Ctr and dsG_VG vs. Ctr comparisons, but not in dsG_V vs. dsG and dsG_VG vs. dsG comparisons (Fig. 5B-E). This suggests that contribution of virus infection itself to the enrichment of genes related to these KEGG terms was not significant in our experimental conditions. Furthermore, the Plant-pathogen interaction and MAPK signaling pathways were still overrepresented in dsG_VG vs. dsG_V comparison (Fig. 5F) albeit with a reduced number of DEGs, suggesting that the interaction of dsGFP with PVX-GFP (RNAi) altered gene expression differentially compared to the interaction of dsGFP with PVX (PTI). Indeed, the overrepresentation of the Plant-pathogen interaction and MAPK signaling pathway terms in the dsG_VG vs. Ctr comparison was greater than in dsG_V vs. Ctr [*p*=6.5E-15 (dsG_VG vs. Ctr) and *p*=8.1E-12 (dsG_V vs. Ctr) for Plant-pathogen interaction; and *p*=2.4E-07 (dsG_VG vs. Ctr) and *p*=2.2E-3 (dsG_V vs. Ctr) for MAPK signaling pathway], indicating a deeper impact of the RNAi response on the transcriptome (Table S2). The KEGG term Plant hormone signal transduction was uniquely overrepresented in the dsG_V vs. Ctr comparison, and includes genes involved in auxin metabolism and signaling, abscisic acid (ABA), cytokinin, gibberellin and ET signaling, and SA response (Table S3).

**Figure 5:**
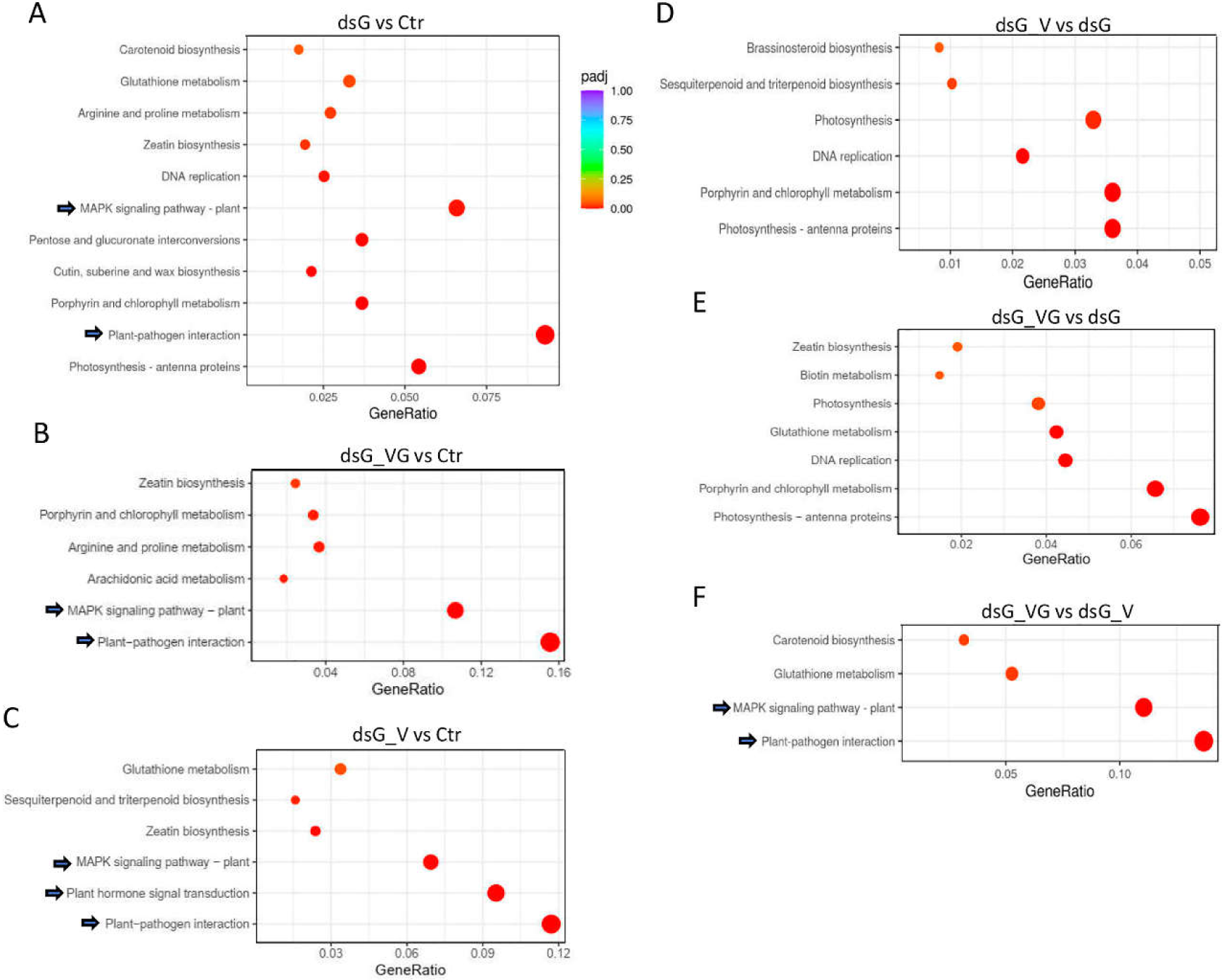
Kyoto Encyclopedia of Genes and Genomes (KEGG) pathway enrichment analysis of differentially expressed genes (DEGs). KEGG terms enriched (adjusted *p*-value < 0.05) in the (A) dsG vs. Ctr, (B) dsG_VG vs. Ctr, (C) dsG_V vs. Ctr, (D) dsG_V vs. dsG, (E) dsG_VG vs. dsG and (F) dsG_VG vs. dsG_V comparisons are shown. Gene ratio is the percentage of total DEGs in the given KEGG term. Dot size represents the number of genes annotated to a specific KEGG term. The KEGG terms Plant-pathogen interaction, MAPK signaling and Plant hormone signal transduction are indicated by arrows.

The dsG vs. Ctr comparison differentially altered the expression of 48 (38 upregulated and 10 downregulated) and 34 (21 upregulated and 13 downregulated) DEGs classified in the Plant-pathogen interaction and MAPK signaling pathways terms, respectively (Table S3). Many of these DEGs were also altered in the datasets from dsG_V vs. Ctr and dsG_VG vs. Ctr comparisons. DEGs in the Plant-pathogen interaction term comprise genes involved in calcium (Ca^+2^) signaling, WRKY transcription factors, acyltransferases, *mitogen-activated protein kinase* (*MAPK*)*3* and *MAPK kinase5*, pathogenesis-related (PR) transcriptional factors, LRR receptor-like serine/threonine-protein kinase *FLS2*, NBS-LRR resistance genes *RPM1* and *RPS2*, EDS1L-like protein, and heat shock protein 82, among others. DEGs in the MAPK signaling pathway term comprise genes involved in ET signaling, Ca^+2^ signaling, WRKY transcription factors, ABA signaling, serine/threonine-protein kinases, among others. A pictorial representation of the non-redundant list of genes associated with the Plant-pathogen interaction and MAPK signaling pathways in the dsG vs. Ctr comparison is showed in Figure 6.

**Figure 6:**
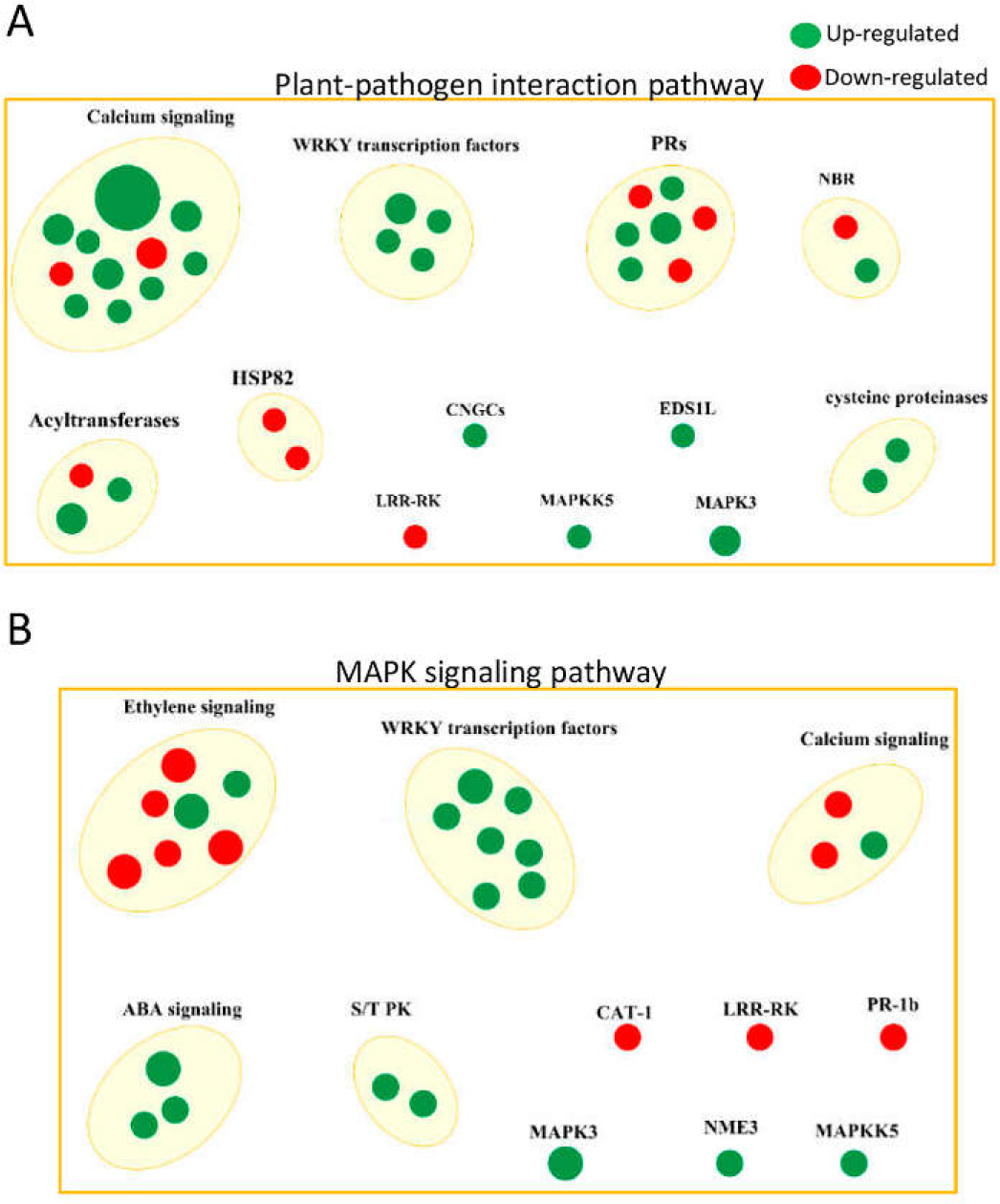
Pictorial representation of the non-redundant list of differentially expressed genes (DEGs) associated with the KEGG term (A) Plant-pathogen interaction and (B) MAPK signaling in the dsG vs. Ctr comparison. DEGs were grouped according to their biological function. Dot size represents the number of DEGs annotated to a specific biological function.

To independently validate the RNAseq results, differential expression of several upregulated genes classified in the Plant-pathogen interaction and MAPK signaling pathways [*MAPK3* (Niben101Scf02171g00008), *WRKY transcription factor6* (*WRKY6*, Niben101Scf02430g03006), *Calcium-binding EF-hand family protein* (*CaEF*, Niben101Scf13289g00008), *1-aminocyclopropane-1-carboxylate synthase2* (*ACS2*, Niben101Scf02334g00004) and the NBS-LRR resistance gene *RPS2* (Niben101Scf10333g00018)] was determined by qRT-PCR using RNA preparations extracted from a new set of samples (not used for RNAseq) derived from dsG, dsG_VG, dsG_V and Ctr treatments (Fig. 7). These candidate genes were selected for their predicted biological functions associated with pathogen interaction, potentially contributing to the RNAi and PTI responses. In general, the gene expression levels measured by qRT-PCR in the different treatments confirmed the pattern observed by RNAseq analyses. While fold-change patterns correlated, discrepancies in magnitude between the qRT-PCR and RNAseq platforms is not uncommon, and could be attributed to differences in the normalization methods used. The relative accumulation of the mRNAs was greater in dsG_VG compared to both dsG_V and dsG treatments. In addition, samples treated with dsG expressed significantly higher levels of ACS2, MAPK3 and CaEF mRNAs compared to controls.

**Figure 7:**
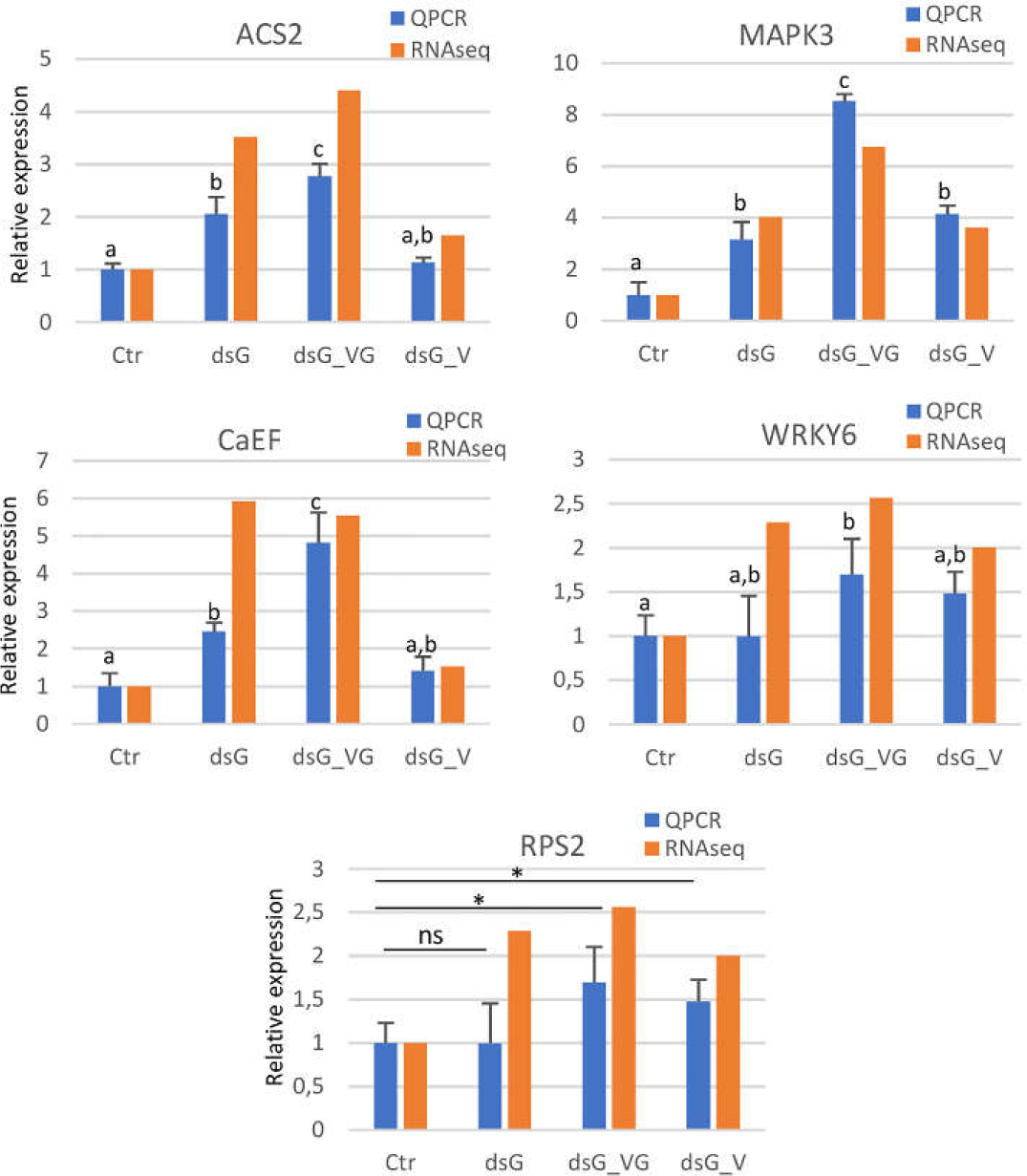
Validation of RNAseq data of representative genes from the KEGG terms Plant-pathogen interaction and MAPK signaling. The relative expression levels of *Mitogen-activated protein kinase3* (*MAPK3*), *WRKY transcription factor6* (*WRKY6*), *Calcium-binding EF-hand family protein* (*CaEF*), *1-aminocyclopropane-1-carboxylate synthase2* (*ACS2*) and the NBS-LRR resistance gene *RPS2* was determined by qRT-PCR using RNA preparations extracted from dsG, dsG_VG, dsG_V and Ctr treatments. Different letters indicate significant differences determined by employing Scheffé’s multiple range test (*p* < 0.05). Asterisks indicate significant differences between treatments (Student’s t-test, *p* < 0.05); ns: not significant. The relative expression levels of selected genes based on the number of Fragments Per Kilobase of transcript sequence per Millions of base-pairs sequenced (FPKM) in each of the three independent samples per treatment used for RNA-Seq are shown for comparison.

### PTI-based defense to PVX-GFP is affected in *AGO2* and *DCL* mutants

*Dicer-like2* (*DCL2*, Niben101Scf08272g00021) and *Argonaute2* (*AGO2*) (Niben101Scf05245g01007) were induced 2.2- and 2.1-fold respectively in the dsG_V vs. Ctr comparison (Table S1). In order to evaluate the role of RNAi genes in PTI induced by dsRNA, we used a *N. benthamiana* transgenic line suppressed for a combination of *DCL2* and *DCL4* (Dcl2/4i) and also a homozygous AGO2*-*/*-* null mutant line (Ago2KO). Wt, Dcl2/4i and Ago2KO plants were inoculated with mixtures of PVX-GFP and either dsGFP, dsPVY or control extracts. dsGFP led to a severe reduction of PVX-GFP RNA levels in the inoculated leaves of Wt plants but not in Dcl2,4i and Ago2KO mutant plants, as assayed by northern blot at 4 dpi (Fig. 8A). Moreover, the accumulation of PVX-GFP RNAs was also diminished in Wt plants treated with dsPVY. However, Dcl2,4i and Ago2KO mutants were insensitive to the PTI response. Comparative analysis by qRT-PCR revealed that the level of PVX-GFP RNA was indeed significantly increased in dsPVY-treated leaves of Dcl2,4i and Ago2KO mutant lines (78 and 27%, respectively) compared to the controls (Fig. 8B). This was corroborated by western blot analysis, where PVX-GFP CP accumulated at greater levels compared to controls in dsPVY-treated Dcl2,4i and Ago2KO plants, and by contrast, less in Wt leaves treated with dsPVY than in control leaves (Fig. 8C).

**Figure 8:**
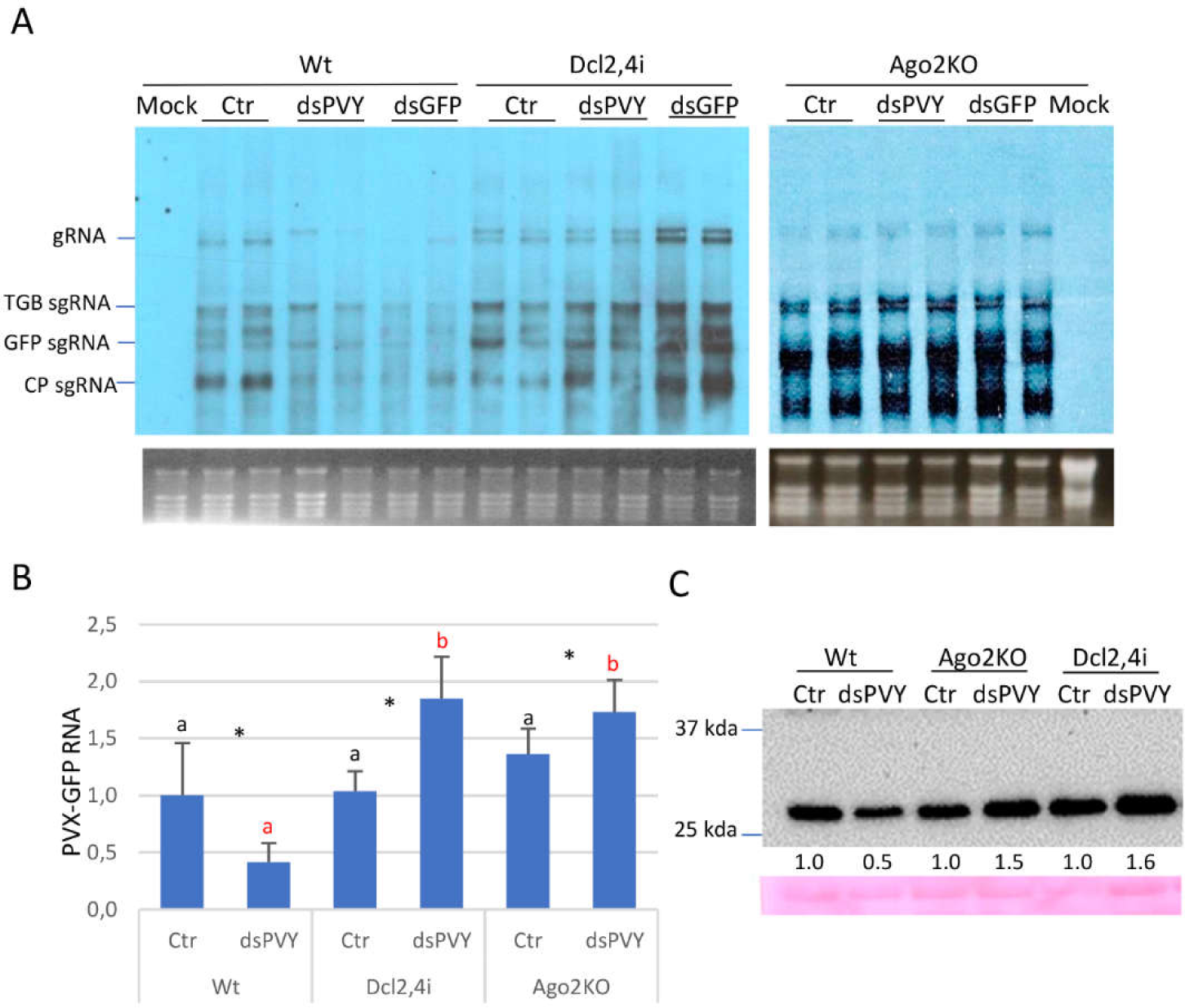
Role of *DCL* and *AGO2* genes in PTI induced by dsRNA. Wt and DCL2/4i and Ago2KO mutant lines were inoculated with mixtures of PVX-GFP and either dsGFP, dsPVY or control (Ctr) extracts. (A) Northern blot analysis of total RNA extracted from inoculated leaf tissues at 4 dpi. Two independent pooled samples were analyzed for each combination. Total RNA (5 µg) was hybridized with a probe complementary to PVX CP. PVX genomic (g)RNA and the major subgenomic (sg)RNAs, triple-gene-block (TGB) sgRNA, GFP sgRNA and CP sgRNA, are indicated. Ethidium bromide staining of rRNA is shown as loading control. (B) qRT-PCR was used to analyze the accumulation of PVX-GFP genomic RNA levels in the inoculated leaves at 4 dpi. Expression of the 18S rRNA gene served as a control. Data represent the means ± SD of 3 replicates, each consisting of a pool of 6 plants that received the same treatment. Different letters indicate significant differences determined by employing Scheffé’s multiple range test for between-group comparisons (*p* < 0.05). For pairwise comparisons, asterisks indicate significant differences between treatments (Student’s t-test, *p* < 0.05); ns: not significant. (C) Western blot analysis of plant extracts derived from inoculated leaves at 4 dpi, using antibodies against PVX CP. The panel below the blot is the membrane stained with ponceau S as control of loading. The intensity of each PVX CP band was quantified by densitometry analyses. Protein levels in control-treated leaves are given the value of 1 and data from samples treated with dsPVY were calculated relative to this value for each genotype. Experiments were repeated once more with similar results.

In independent experiments, we introduced *N. benthamiana* transgenic lines suppressed for either DCL2 (Dcl2i) or DCL4 (Dcl4i), together with Wt and Dcl2,4i plants in the analysis. At 4 dpi, there was a reduction in the number of GFP foci in the leaves of Wt and Dcl2i plants inoculated with PVX-GFP plus dsGFP compared to controls, but not in Dcl4i and Dcl2,4i plants (Fig. 9A). By contrast, there was no significant difference in the number of GFP foci between dsPVY- and control-treated leaves in all the genotypes analyzed. Western blot analyses from a pool of inoculated leaves (two replicates) revealed that PVX-GFP in combination with dsPVY accumulated at greater levels in transgenic lines suppressed for *DCL* genes compared to controls, in particular in Dcl2i plants (Fig. 9B). Furthermore, western blot analysis from a pool of upper leaf tissues from each treatment (two replicates) at 7 dpi revealed that accumulation of PVX-GFP CP was lower in Wt plants treated with PVX-GFP plus dsPVY than in control-treated plants, whereas PVX-GFP in combination with dsPVY accumulated at greater levels compared to controls in Dcl2,4i and Dcl2i plants (Fig. 9C). PVX-GFP accumulated at roughly similar levels in the systemic leaves of Dcl4i plants treated with either dsPVY or control extract. Taken together, our findings suggest that AGO2, DCL2 and DCL4 seem to be positively involved in PTI-based defense against virus infection

**Figure 9:**
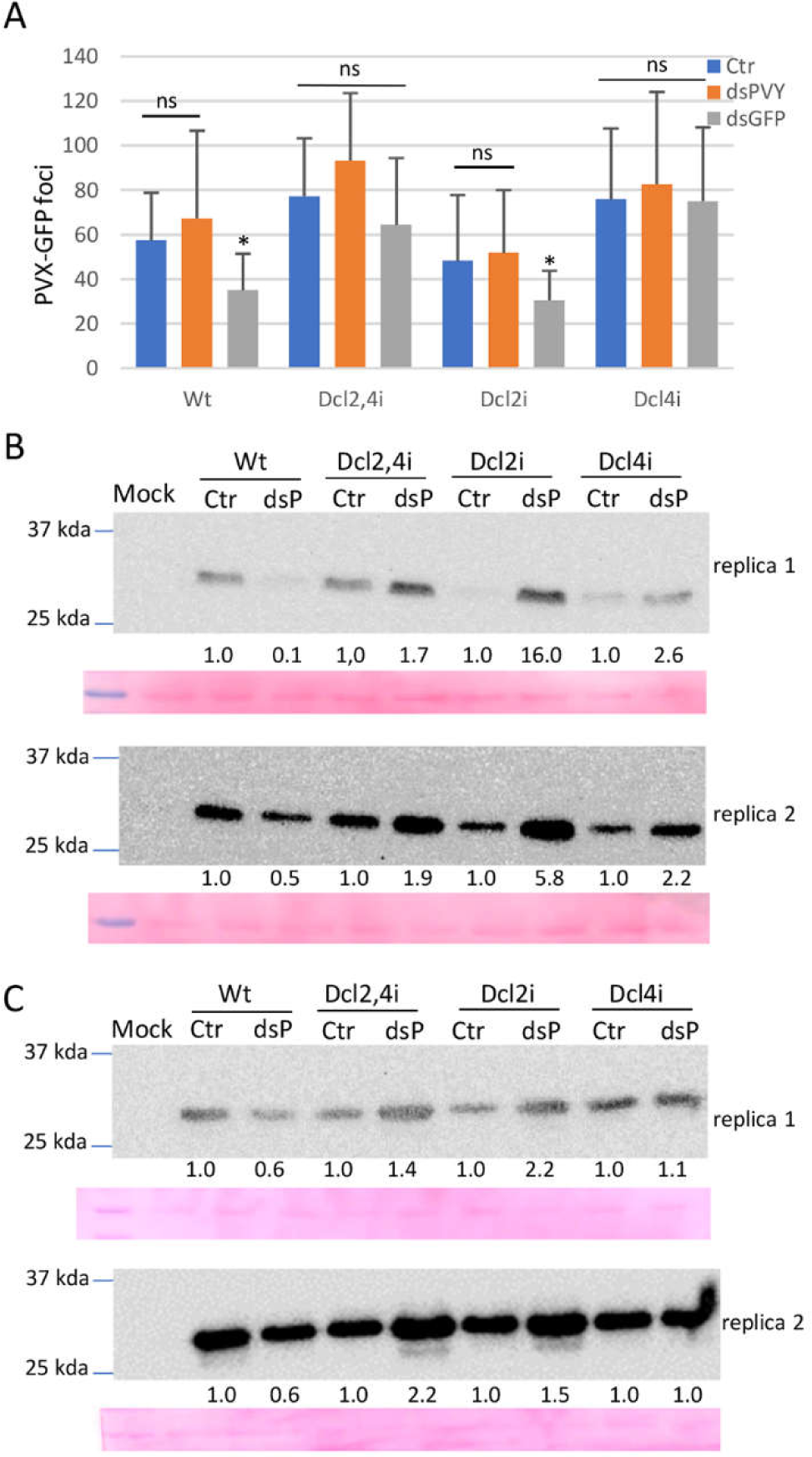
Local and systemic responses to dsRNA in single and double DCL mutant lines. Wt and DCL2i, DCL4i, and DCL2/4i mutant lines were inoculated with mixtures of PVX-GFP and either dsGFP, dsPVY or control (Ctr) extracts. (A) Mean numbers ± standard deviation of infection foci on inoculated leaves of 6 plants (two leaves per plant) with the different treatments at 4 days post-inoculation (dpi). Asterisks indicate significant differences between treatments (Student’s t-test, *p* < 0.05); ns: not significant. (B) Western blot analysis of plant extracts derived from inoculated leaves at 4 dpi, using antibodies against PVX CP. Independent pooled samples were analyzed in two replica assays. dsP: dsPVY (C) Western blot analysis of plant extracts derived from systemic leaves at 7 dpi, using antibodies against PVX CP. The panels below the blots are the membranes stained with ponceau S as control of loading. The intensity of each PVX CP band was quantified by densitometry analyses. Protein levels in control-treated leaves are given the value of 1 and data from samples treated with dsPVY were calculated relative to this value for each genotype. Experiments were repeated once more with similar results.

### PTI-based defense to PVX-GFP is enhanced by SA signaling

In order to determine whether signaling by the two major phytohormones involved in defense contributed to the PTI response against PVX-GFP, *N. benthamiana* transgenic plants deficient in JA perception (*COI1* IR line) or SA accumulation (*NahG* line), as well as the Wt plants, were inoculated with mixtures of PVX-GFP plus either dsPVY or control extracts. At 4 dpi, there were no significant differences in the number of GFP foci between the leaves treated with dsPVY and control-treated leaves in *COI1* IR and Wt plants (Fig. 10A). However, there was a greater number of GFP foci in the inoculated leaves treated with dsPVY than with the control in *NahG* transgenic plants. Both viral RNAs and CP accumulated less in the inoculated leaves treated with PVX-GFP plus dsPVY than in control-treated leaves in both *COI1* IR and Wt plants, whereas PVX-GFP accumulated at greater levels in the treatment with dsPVY compared to control in *NahG* plants as assayed by northern and western blot analyses (Fig. 10B,C).

**Figure 10:**
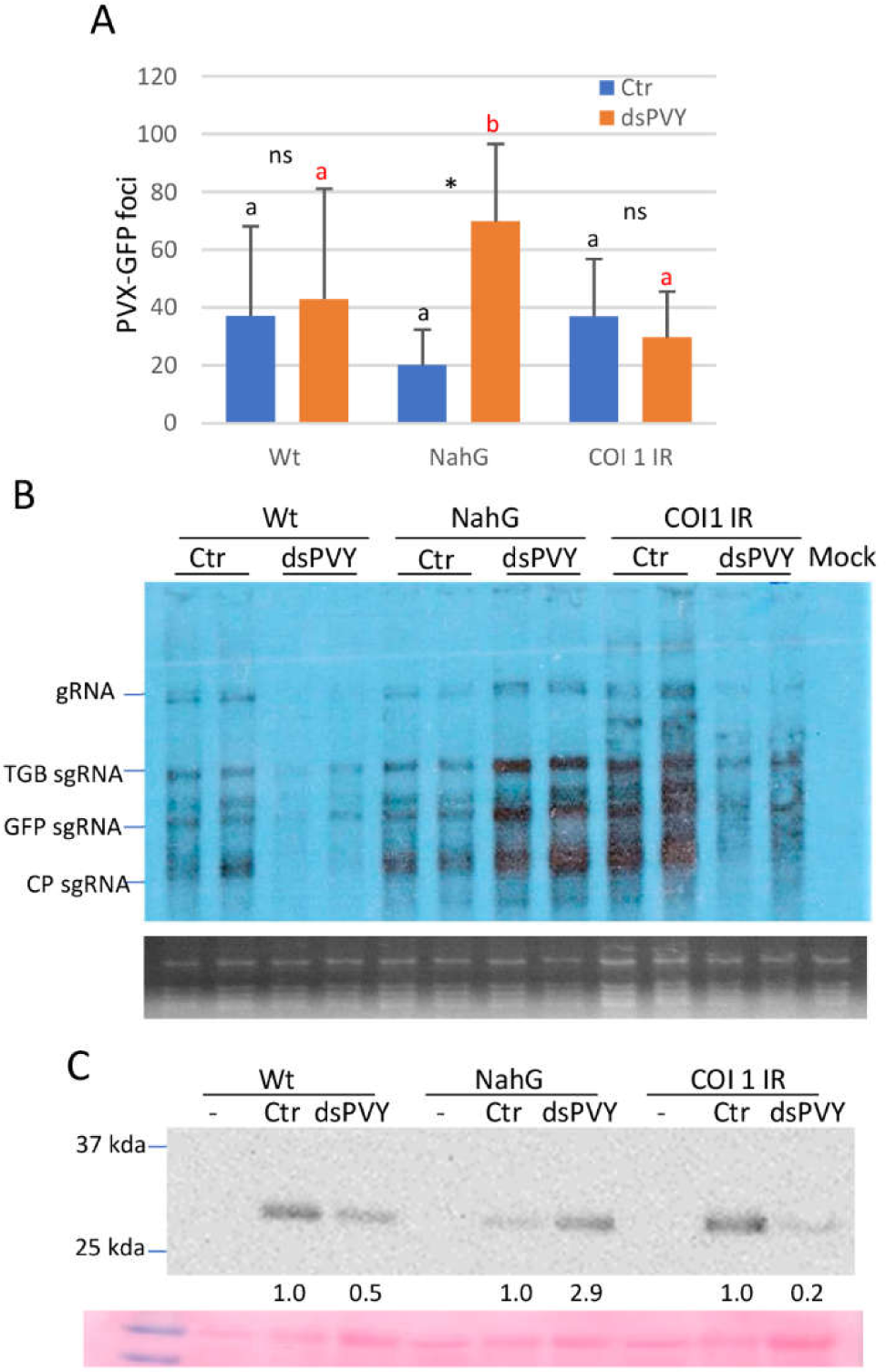
PTI response is affected by SA signaling pathway. *Nicotiana benthamiana* transgenic plants deficient in jasmonic acid perception (*COI1* IR) or salicylic acid accumulation (*NahG*), as well as the Wt plants, were inoculated with mixtures of PVX-GFP plus either dsPVY or control extracts. (A) Mean numbers ± standard deviation of infection foci on inoculated leaves of 9 plants (two leaves per plant) with the different treatments at 4 days post-inoculation (dpi). Different letters indicate significant differences determined by employing Scheffé’s multiple range test for between-group comparisons (*p* < 0.05). For pairwise comparisons, asterisks indicate significant differences between treatments (Student’s t-test, *p* < 0.05); ns: not significant. (B) Northern blot analysis of total RNA extracted from inoculated leaf tissues at 4 dpi. Two independent pooled samples were analyzed for each combination. Total RNA (5 µg) was hybridized with a probe complementary to PVX CP. PVX genomic (g)RNA and the major subgenomic (sg)RNAs, triple-gene-block (TGB) sgRNA, GFP sgRNA and CP sgRNA, are indicated. Ethidium bromide staining of rRNA is shown as loading control. (C) Western blot analysis of plant extracts derived from inoculated leaves at 4 dpi, using antibodies against PVX CP. Non-infected plants from each genotype were used as negative controls (-). The panel below the blot is the membrane stained with ponceau S as control of loading. The intensity of each PVX CP band was quantified by densitometry analyses. Protein levels in control-treated leaves are given the value of 1 and data from samples treated with dsPVY were calculated relative to this value for each genotype. The experiments were repeated twice with similar results.

To further test whether the SA signaling pathway affected the PTI response, *N. benthamiana* Wt plants were treated with SA, methyl JA (MeJA) or mock solution and then inoculated with PVX-GFP plus either dsPVY or control extracts. At 4 dpi, there was a significant decrease in the number of GFP foci in SA-treated leaves inoculated with dsPVY, whereas treatment with MeJA induced an increase in the number of PVX-GFP-derived foci compared to controls (Fig. 11A). Western blot analysis revealed that PVX CP accumulated less in leaves inoculated with PVX-GFP plus dsPVY than in control leaves in both mock and SA-treated plants (Fig. 11B). By contrast, PVX-GFP in combination with dsPVY accumulated at greater levels compared to control in plants treated with MeJA. As expected, SA treatment induced the expression of the SA-responsive gene *Pathogenesis-related protein-1* (*PR-1*), in particular in dsPVY-inoculated leaves, as assayed by qRT-PCR (Fig. 11C). No significant difference in PR-1 mRNA levels was found in virus infected-plants treated with MeJA as compared to non-infected plants treated with mock solution. Altogether, these findings suggest that dsRNA-induced PTI is enhanced by SA signaling and that the effect of MeJA in the enhanced accumulation of PVX-GFP in dsPVY-treated plants depends on the antagonistic relationship between JA and SA signaling pathways.

**Figure 11:**
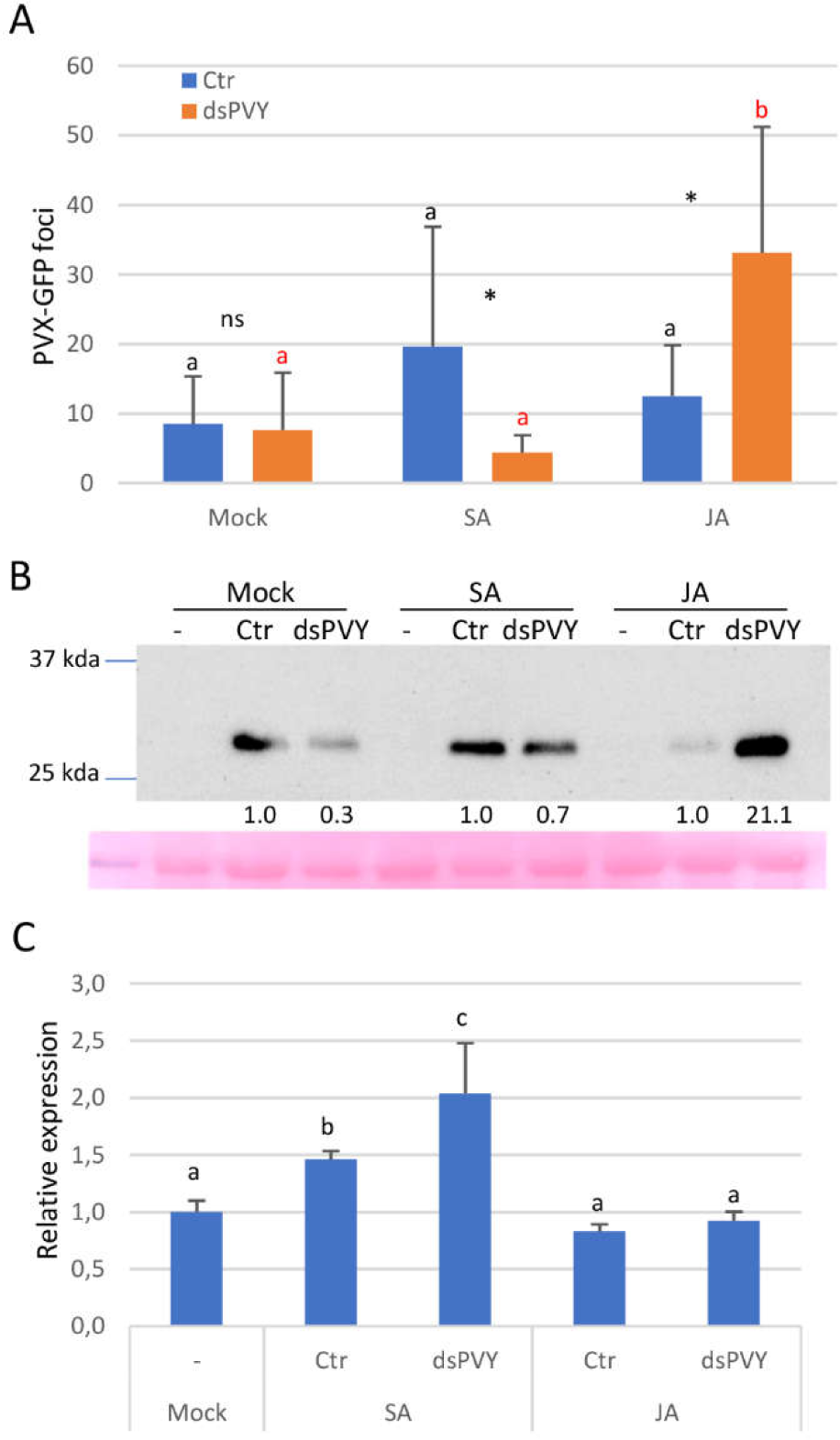
Effects of hormone treatments in the PTI response against PVX-GFP. Salicylic acid (SA), methylJA (JA), or mock solution as a control, was applied to wild-type *Nicotiana benthamiana* plants that were mock-inoculated or inoculated with mixtures of PVX-GFP plus either dsPVY or control extracts. (A) Mean numbers ± standard deviation of infection foci on inoculated leaves of 8 plants (two leaves per plant) with the different treatments at 4 days post-inoculation (dpi). Different letters indicate significant differences determined by employing Scheffé’s multiple range test for between-group comparisons (*p* < 0.05). For pairwise comparisons, asterisks indicate significant differences between treatments (Student’s t-test, *p* < 0.05); ns: not significant. (B) Western blot analysis of plant extracts derived from inoculated leaves at 4 dpi, using antibodies against PVX CP. Non-infected plants from each genotype were used as negative controls (-). The panel below the blot is the membrane stained with ponceau S as control of loading. The intensity of each PVX CP band was quantified by densitometry analyses. Protein levels in control-treated leaves are given the value of 1 and data from samples treated with dsPVY were calculated relative to this value for each hormone treatment. (C) qRT-PCR was used to analyze the accumulation of Pathogenesis-related protein-1 RNA levels in the inoculated leaves at 4 dpi. Expression of the 18S rRNA gene served as a control. Data represent the means ± SD of 3 replicates, each consisting of a pool of 8 plants that received the same treatment. Statistical comparisons between means were made by employing Mann-Whitney U test with a Bonferroni correction for multiple comparisons of α to α= 0.005. Different letters indicate significant differences at *p* < 0.005. The experiments were repeated once more with similar results.

## DISCUSSION

One of the main goals of this study was to compare the differential efficiency of the antiviral activity triggered by externally delivered, virus-specific and nonspecific dsRNAs in plants. RNAi mediated by virus-specific dsRNA is the main antiviral defense mechanism in plants, but nonspecific dsRNA-triggered responses have been documented to play a role in antiviral defense (13, 17). Using PVX-GFP as a model, we have determined that co-inoculation with either virus-specific or nonspecific dsRNA reduced virus accumulation in both inoculated and systemic leaves, although at different extents (Fig. 1 and Fig. 2). While the administration of dsRNA specific for the targeted virus induced a potent RNAi-based antiviral response that resulted in highly effective control of viral disease, the degree of interference with PVX-GFP infection afforded by nonspecific dsRNA (PTI) was limited; viral titers in the leaves treated with nonspecific dsRNA were several-fold higher than in the leaves treated with virus-specific dsRNA, and the plants became susceptible to systemic infection. Thus, our data point out that nonspecific dsRNA is a poor inducer of antiviral immunity compared to virus-specific dsRNA capable to trigger the RNAi response. The coexistence of non-sequence-specific immunity and RNAi as two distinct antiviral mechanisms induced by dsRNA has also been reported in invertebrates, albeit nonspecific dsRNA was shown to evoke an antiviral response much lower in potency than that induced by virus-specific dsRNA (12, 28).

It has been reported that nonspecific dsRNA-induced immunity against TMV mainly restricted the cell-to-cell movement of the virus by callose deposition at PD, but did not affect TMV replication (17). Deposition of callose at PD regulates symplastic transport limiting the ability of the invading pathogen to spread throughout the plant (29, 30). We show here that cell-to-cell movement of PVX-GFP was not significantly inhibited by nonspecific dsRNA, as judged by the unaltered size of PVX-GFP infection sites in dsPVY-treated leaves (Fig. 3). Remarkably, studies with a frameshift mutant in the P25 movement protein of PVX indicated that nonspecific dsRNA caused a significant reduction in viral replication. PVX mutants defective in the P25 protein do not move out of the initially infected cell, although they do accumulate at Wt levels in individual cells (27,31). Overall, our data support a model where interference with PVX-GFP accumulation triggered by nonspecific dsRNA operates at the single-cell level, and lead to reduce accumulation of the virus in the inoculated leaves. Thus, effects of PTI triggered by dsRNA on virus infection may vary with the specific virus-host combination. Additionally, it cannot be excluded that proteins encoded by PVX-GFP act as effectors to suppress dsRNA-induced immunity based on restriction of viral cell-to-cell movement, as has been reported previously for other viral proteins (17, 18). The bacterial PTI elicitor derived from flagellin, flg22, is known to trigger deposition of callose at PD and restricts TMV cell-to cell movement in *N. benthamiana* (17). Our results showed that PD callose deposition elicited by flg22 was correlated with a diminished accumulation of PVX-GFP in the inoculated leaves but did not affect virus replication, suggesting that different PTI elicitors, i.e., dsRNA and flg22, may restrict PVX-GFP at distinct steps of virus infection. In this sense, unlike for dsRNA, PTI triggered by flg22 has been shown to induce the production of ROS species in *N. benthamiana* and *A. thaliana* (17).

Previous findings showed that PVX infection was not affected by nonspecific dsRNA in spite of the induction of PTI-like responses (14). However, careful examination of the experimental conditions revealed differences between the work presented here, which examined the effect of co-inoculation of dsRNA on the accumulation of the virus in the inoculated leaves, and the other study, where potato plants were challenged with PVX 24 h after dsRNA application and accumulation of PVX RNA was examined in systemically infected leaves. Our observation that accumulation of PVX-GFP in systemically infected leaves of symptomatic plants was only slightly diminished by nonspecific dsRNAs is consistent with this discrepancy. In a broader sense, our findings could explain other examples in the literature where dsRNA did not confer systemic antiviral protection against heterologous viruses (32, 33).

Before this study, no attempt had been reported to examine the whole transcriptomic response to dsRNA either alone or in the context of infections with homologous and heterologous viruses to shed light on differences between these responses. It is noteworthy that the experimental approach implemented in this work, with samples taken at an early stage of local infection, allowed us to investigate the outputs associated with early events in dsRNA signaling in the absence of the robust transcriptomic response triggered by PVX at later stages of infection (34). PCA analysis revealed that treatment with dsRNA alone had an ample effect on host gene expression, which could be responsible in part for the antiviral responses triggered by dsRNA (Fig. 4). KEGG analysis showed a significant enrichment of terms related to plant-pathogen signaling pathways (KEGG terms Plant-pathogen interaction and MAPK signaling) in all the three treatments that included dsRNA (Fig. 5). Our results further indicated that the transcriptomic response triggered by dsRNA alone included canonical immune pathways or genes known to be involved in defense responses, i.e., Ca^+2^ signaling, ET signaling, MAPK signaling, WRKY transcription factors, PR transcriptional factors, NBS-LRR resistance genes, EDS1, and LRR receptor-like kinases, many of which are typical of antimicrobial PTI (Fig. 6) (11, 13, 35, 36). Moreover, the transcriptomic response to the homologous combination (dsGFP plus PVX-GFP) had a greater overrepresentation of genes involved in plant-pathogen signaling pathways than the heterologous combination (dsGFP plus Wt PVX), highlighting quantitative differences between RNAi and PTI immune responses (Table S2). These findings are in accordance with the differential efficiency in antiviral immunity triggered by virus-specific vs. nonspecific dsRNA. Moreover, this observation is somehow reminiscent to the intimate relationships between PTI and effector-triggered immunity (ETI) (11, 37). Although these two types of immune pathways involve different activation circuits, their downstream outputs (Ca^+2^ flux, ROS burst, MAPK cascades, transcriptional reprograming and phytohormone signaling) usually converge, albeit with differences in amplitude and duration.

It is noteworthy to mention that Ca^+2^ signaling has been involved in antiviral immunity against TMV triggered by nonspecific dsRNA (17). Furthermore, it has been reported that a wound-induced, Ca^2+^ signaling cascade stabilizes mRNAs encoding key components of RNAi machinery, notably AGO1/2, DCL1 and RNA-dependent RNA polymerase6, enhancing plant defense against virus infection (38). Thus, it is tempting to speculate that the strong Ca^+2^ signaling evoked by dsRNA in this study activated RNAi-related gene expression and resulted in highly effective control of viral disease when the dsRNA shares sequence homology with the cognate virus. This was somehow corroborated in our transcriptomic analysis, where a significant induction of AGO2 and DCL2 mRNAs was detected by RNAseq analysis (Table S1).

AGO2, DCL2 and DCL4 play a moderate role, if any, in restricting the accumulation of PVX-GFP in local infections in *N. benthamiana* (7, 10, 39). Concordantly, accumulation of PVX-GFP RNA in control-treated, local leaves of Ago2KO and Dcl2,4i plants did not vary significantly compared to Wt plants (Fig. 8). However, treatment with virus-specific dsRNA in Wt and Dcl2i plants but not in Ago2KO, Dcl4i and Dcl2,4i plants interfered with local infection of PVX-GFP, indicating that AGO2 and DCL4 participate in resistance to virus infection conferred by homologous dsRNA. Even more remarkable is that down-regulation of AGO2, and DCL2 and DCL4 in single and double mutants abrogated antiviral resistance conferred by nonspecific dsRNA, suggesting a connection between RNAi-related genes and dsRNA-induced PTI defense (Fig. 8 and Fig. 9). Previously, it was reported that nonspecific dsRNA-induced PTI responses, i.e., MAPK activation and ET production, were not impaired in Arabidopsis mutants deficient in DCL2 and DCL4 (13). However, interference with virus accumulation triggered by nonspecific dsRNA in these plants was not addressed. Non-canonical roles of RNAi-related genes in antiviral defense mechanisms have been previously documented. In *Caenorhabditis elegans*, the Dicer-Related Helicase DRH1 acts as a PRR in response to virus infection, and activates an immune response associated with increased pathogen resistance (40). In addition to its involvement in RNAi, the *Drosophila* Dicer-2 mediates defense gene expression in a tissue- and virus-specific manner (41). Furthermore, recent findings in Arabidopsis have revealed a function of DCL2 in activation of basal antiviral immunity in response to dsRNA through NBS-LRR proteins, which was independent of its canonical function in RNAi (42). Regarding Argonaute proteins, AGO4 has been implicated in the degradation-independent translational control of PVX-GFP transcripts upon elicitation of NBS-LRR-mediated resistance (43). Moreover, the tethering of human AGO2 and AGO4 proteins to mRNAs via fusion proteins was sufficient to inhibit translation in a sRNA-independent manner (44). Although somewhat speculative, it is conceivable that some of the viral susceptibility in AGO2, DCL2 and DCL4 mutant plants that has been attributed to defects in RNAi could instead be due to defects in PTI activation. At present, it is unknown why nonspecific dsRNA stimulates rather than suppresses PVX-GFP accumulation in AGO2 and DCL mutant genotypes. In addition to AGO2, antiviral activities associated with AGO5L and AGO7 have been described in *N. benthamiana* (45). Moreover, AGO5 has been shown to act synergistically together with AGO2 to restrict PVX accumulation (46). Similarly, although DCL4 and DCL2 are the main contributors of antiviral RNA silencing, DCL3 has a minor but significant role in defense against several viruses (6, 47). Thus, we could hypothesize that nonspecific dsRNA might be saturating the remaining RNA silencing machinery presents in Ago2KO and DCL2i/4i mutants, rendering these plants more susceptible to PVX-GFP accumulation. Nonspecific dsRNA may be recognized and cleaved by DCL proteins and hence may outcompete with viral replication intermediates for access to the RNAi machinery. Further work would be needed to validate this hypothesis and the involvement of RNAi-related genes in dsRNA-induced PTI defense.

The altered expression of genes classified in the KEGG term Plant hormone signal transduction detected by RNAseq analysis suggested an important contribution of hormone signaling in the PTI response to virus infection. There are precedents arguing for a role of the phytohormones SA and JA in PTI-based defenses induced by exogenous application of bacterial RNA derived from *P. syringae* in *Arabidopsis* (21). In addition, SA was required for both local and systemic resistance induced by bacterial PAMPs like flg22 and lipopolysaccharides against *P. syringae* (48). Arabidopsis mutants affected in SA biosynthesis and signaling were impaired in establishing the systemic resistance response triggered by bacterial PAMPs. In our study, a role for SA signaling in the establishment of PTI induced by nonspecific dsRNA in virus-infected plants was substantiated from the analysis of SA-deficient transgenic plants and treatments with phytohormones (Fig. 10 and Fig. 11). Preventing SA accumulation in plants expressing the *NahG* transgene increased the number of GFP foci derived from PVX-GFP and virus accumulated at higher levels in leaves treated with dsRNA compared to control-treated leaves. Conversely, treatment with SA enhanced the inhibitory effect of nonspecific dsRNA on PVX-GFP infectivity, as reflected by the reduced number of GFP foci. This was correlated with an increased expression of the SA-responsive gene *PR-1* in plants treated with dsRNA compared to controls. Furthermore, the enhanced virulence of PVX-GFP in *NahG* plants was mimicked by treatment of Wt plants with MeJA, a plant hormone known to inhibit or down-regulate SA-mediated defenses (49, 50). It has been reported that the application of exogenous SA suppresses the JA signaling pathway (51). Thus, it could be argued that the effect of SA on PTI induced by dsRNA might be mediated by interference with JA-dependent defense gene expression. However, the inhibitory effect of nonspecific dsRNA on PVX-GFP infectivity seemed to be independent from JA-mediated signaling, as the transgenic plants insensitive to JA (*COI1* IR) showed susceptibility to PVX-GFP similar to Wt plants. *COI1* participates in JA perception and regulates almost all the metabolic processes responding to JA (52). In addition to SA, we identified several DEGs involved in other hormone signaling pathways in the dataset from dsRNA-treated plants, suggesting that other phytohormones might interact with SA in the PTI response to dsRNA (Table S3). In this scenario, ET enhanced SA-responsive *PR-1* expression in Arabidopsis, and in tobacco it was crucial for the onset of systemic acquired resistance (53, 54).

In summary, we show here that although PTI-based defenses triggered by nonspecific dsRNA reduced virus accumulation in both local and systemic tissues, antiviral immunity conferred by nonspecific dsRNA was weaker compared to the RNAi response triggered by virus-specific dsRNA. Such differential efficiency in virus reduction was correlated with a deeper impact on defense responses in plants treated with the homologous dsRNA-virus combination compared to those elicited by the heterologous dsRNA-virus combination. Further, we demonstrated that unlike other examples of dsRNA-based PTI, which restricted TMV cell-to cell movement, PTI induced by nonspecific dsRNA partially inhibited PVX-GFP accumulation at the single-cell level. Our data also suggested an unexpected connection between RNAi-related genes and dsRNA-induced PTI defense, which needs to be further investigated.

## MATERIALS AND METHODS

### Plant materials

The *N. benthamiana* transgenic plants expressing the *Salicylate hydroxylase* gene (*NahG*) and the *COI1* IR line, in which *NbCOI1* was silenced by an RNAi hairpin construct, have been described previously (55, 56). *N*. *benthamiana* plants expressing hairpins that target endogenous DCL2, DCL4 and DCL2,4 transcripts were previously described (57). In addition, a knockout mutant of *AGO2* generated by CRISPR/Cas9 technology was used (10). Plants were kept in environment-controlled growth chambers with 16/8 h day/night photoperiod, ∼2500 lux of daylight intensity and 60% relative humidity.

### Plasmid construct

The complete CP coding sequence and flanking regions of PVY (902 bp) was cloned into L4440 as described (58). L4440 is a plasmid vector which has two convergent T7 promoters flanking the multiple cloning sites (59). The complete GFP coding sequence (717 bp) was amplified by PCR using pSLJ-GFP (60), and cloned into *Sac*I and *Pst*I of L4440. The upstream primer was 5’ *GAGCTC*ATGGCAAGTAAAGGAGAAGAAC 3’ (italized sequence corresponds to the *Sac*I restriction site). The downstream primer was 5’ *CTGCAG*TTTGTATAGTTCATCCATGCCAT3’ (italized sequence corresponds to the *Pst*I restriction site). Plasmids were transformed into *Escherichia coli* HT115(DE3) using standard CaCl_2_ transformation protocols. HT115(DE3) is an RNAase III-deficient *E. coli* strain, which was modified to express T7 RNA polymerase from an isopropyl β-D-1-thiogalactopyranoside (IPTG)-inducible promoter (59).

The binary vector pGR107 expressing the infectious cDNA of PVX-GFP has been previously described (61). To construct PVXΔMP-GFP, pGR107 was digested with *Apa*I (nucleotide 4945 of PVX), end filled with Klenow fragment, and religated (62). The PVX-GFP binary vector was previously described (63).

### dsRNA production

Single colonies of HT115(DE3) containing the L4440 plasmid derivatives were grown at 37**°**C for 16 h in Luria-Bertani (LB) broth with ampicillin and tetracycline at a final concentration of 500 and 12.5µg/ml, respectively. The culture was diluted 75-fold in the same medium and allowed to grow to OD_595_= 0.5. T7 polymerase was induced by the addition of 10 µM IPTG, and the culture was incubated further with shaking for 2 h at 37**°**C. After that, the culture broths were centrifuged (4,000×rpm, 15 min) to collect the cells, and bacterial pellets were re-suspended in 1 M ammonium acetate (64). Total nucleic acid was extracted after a phenol-chloroform step prior to ethanol precipitation. The nucleic acids prepared using non-induced cultures of HT115(DE3) containing the L4440 derivatives were used as negative control in all the experiments. Concentration of dsRNA in different preparations was estimated to be approximately 60 ng/µl, as judged by comparison with dsDNA markers.

### Virus inoculation and topical application of dsRNA

In order to ensure the uniformity of the viral inocula in all the experiments, inoculum stocks were prepared from local *N. benthamiana* leaves agro-infiltrated with either Wt PVX, PVX-GFP or PVXΔMP-GFP. For this, symptomatic leaf tissue was cut in small slices (1 cm^2^), split into 500 mg aliquots and stored at -80°C until use. Virus inoculations on 3-to-4-week old plantlets was performed by grinding each aliquot in sodium phosphate buffer (0.02 M, pH 7) at 1:5 (w/v). A 20 µl-dose of infected sap combined with dsRNA extract (1:1 sap:dsRNA ratio; 1.2 µg dsRNA) was applied to two leaves of each plant previously dusted with carborundum as abrasive (Carlo Erba, Barcelona, Spain). The number of PVX-GFP-derived foci was assessed with a Black Ray® long wave UV lamp (UVP, Upland, CA, USA). For GFP foci measurements, 6 leaves from 3 plants for each treatment were scanned with image-analysis software ImageJ (http://imagej.nih.gov/ij/index.htmls).

### RNA and Protein Gel Blot Analysis

To minimize the effects of interleaf variability, leaf tissue from different plants corresponding to the same treatment type were pooled. Total RNA was extracted from inoculated leaves at 4 and 7 dpi and from upper leaves 7 dpi as described (56). RNA samples were separated on 1% agarose formaldehyde gels and transferred to Hybond-N membranes (Roche Molecular Biochemicals). Membrane hybridization was carried out overnight at 65°C using digoxigenin-labeled riboprobes corresponding to PVX CP sequences.

Total proteins were extracted by grounding leaf disks with a pestle and mixed with five volumes of extraction buffer (0.1 M Tris-HCl PH 8, 10 mM EDTA, 0.1 M LiCl, 1% β-mercaptoethanol and 1% SDS) (65). Samples were boiled and fractionated in 15% SDS-PAGE gels. PVX CP was detected with a commercial rabbit antibody (1:1000 dilution) (No. 070375/500; Loewe Biochemica GmbH, Germany) using an appropriate secondary antibody conjugated with horseradish peroxidase. Detection was performed using the ECL system (Amersham Biosciences). Densitometric analysis of blotted protein bands was performed using the public domain software ImageJ (v1.52p) (National Institutes of Health website, image processing and analysis in java).

### qRT-PCR analysis

qRT-PCR for the analysis of gene expression was performed with gene-specific primers (Table S3). The relative quantification of PCR products was calculated by the comparative cycle threshold (ΔΔCt) method as described (56). Virus detection was performed using primers that amplify a region from nucleotides 2621 to 2753 of the PVX sequence. Amplification of 18S rRNA was chosen for normalization because of its similar level of expression across all treatments. All qRT-PCR experiments were performed in triplicate.

### Callose staining

Callose staining was carried out as described (66). Briefly, individual leaf disks were soaked with 0,1% aniline blue solution (in 50 mM potassium phosphate buffer, pH 8.0) containing either 1 μM flg22 (MedChemExpress, NJ, USA), 30 ng/ul dsPVY or control extracts. Leaf disks were further evacuated for 1-2 minutes (< 0.8 Pa) in a vacuum desiccator. Aniline blue fluorescence was imaged 30 minutes after dsPVY/flg22 or control treatment using a Leica TCS SP8 STED 3X confocal microscope (Wetzlar, Germany) with Leica Application Suite X software and using a 405 nm diode laser for excitation and filtering the emission at 430-490 nm. 8-bit images were acquired with a HC PL APO 40x/1.30 Oil CS2 objective. Callose fluorescence intensity was quantified with ImageJ software (http://rsbweb.nih.gov/ij/) using the plug-in calloseQuant (66). Callose spots were measured in 5-6 images taken from three leaf discs per plant from two different plants for each treatment. Regions of interest selected by calloseQuant were verified visually before measurement. ROIs that were not overlapping with the cell wall or did not contain clear signal above the background were deleted. Individual fluorescence intensities that occurred as outliers from the general distribution of fluorescence intensities (<1%) in the sample were excluded from analysis.

### Library Preparation for Transcriptome Sequencing

Three independent biological replicates were used to monitor differences in gene expression between treatments. The integrity and quality of the total RNA were checked using NanoDrop 2000 Spectrophotometer (Thermo Scientific, Waltham, MA, USA) and formaldehyde agarose gel electrophoresis. The transcriptome libraries, sequencing and bioinformatics analysis were performed at Novogene in United Kingdom; mRNA was purified from total RNA using poly-T oligo-attached magnetic beads. After fragmentation, the first strand cDNA was synthesized using random hexamer primers, followed by the second strand cDNA synthesis. The library was checked with Qubit and real-time PCR (Rotor-Gene Q thermal cycler, Qiagen, Hilden, Germany) for quantification and bioanalyzer for size distribution detection. Quantified libraries were pooled and sequenced on Illumina NovaSeq PE150 Platform according to effective library concentration and data amount. The clustering of the index-coded samples was performed according to the manufacturer’s instructions. After cluster generation, the library preparations were sequenced and paired-end reads were generated.

Raw reads were firstly processed through in-house Perl scripts. In this step, clean reads were obtained by removing reads containing adapter, reads containing ploy-N and low-quality reads from raw data. At the same time, Q30 and GC content of the clean data were calculated. All the downstream analyses were based on the clean data with high quality.

The *N. benthamiana* reference genome and gene model annotation files were downloaded from the SGN ftp site (ftp://ftp.solgenomics.net/genomes/Nicotiana_benthamiana/assemblies directly. An index of the reference genome was built using Hisat2 v2.0.5 and paired-end clean reads were aligned to the reference genome using Hisat2 v2.0.5. FeatureCounts v1.5.0-p3 was used to count the reads numbers mapped to each gene. The expected number of Fragments Per Kilobase of transcript sequence per Millions of base-pairs sequenced (FPKM) of each gene was calculated based on the length of the gene and reads count mapped to this gene.

### KEGG Pathway Enrichment Analysis of Differentially Expressed Genes

Differential expression analysis of the two assayed treatments (three replicates per treatment) was performed using the DESeq2 R package (1.20.0) (67). The resulting *p*-values were adjusted using the Benjamini and Hochberg’s correction for controlling the false discovery rate. Genes with a |log2(Fold-change)| ≥ 1 and adjusted *p-*value ≤ 0.05 were assigned as differentially expressed. Fold-change calculations were performed for paired-comparisons made between treatments.

KEGG (Kyoto Encyclopedia of Genes and Genomes) is a collection of manually curated databases containing resources on genomic, biological-pathway and disease information (68). KEGG pathway enrichment analysis identifies significantly enriched metabolic pathways or signal transduction pathways associated with differentially expressed genes, comparing the whole genome background. Enrichment analysis of DEGs was implemented by the clusterProfiler R package (v3.8.1), in which gene length bias was corrected. KEGG pathways with adjusted *p* value less than 0.05 were considered significantly enriched. Gene ontology (GO) enrichment analysis rendered highly generic GO terms and was not used for subsequent analyses.

### Hormone treatments

Four-week-old *N. benthamiana* plants were sprayed with MeJA (50 μM), SA (1 mM) (Sigma) or mock solution (containing 1 mM ethanol) every day from 3 days before viral inoculation as indicated above.

### Statistical Analysis

All statistical analyses were performed using the statistical software IBM SPSS Statistics v.25 (IBM Corp). For each experiment, samples were assessed for normality via the Shapiro–Wilk test and for equality of variances using Levene’s test. For experiments with normally distributed samples of equal variance, one-way ANOVA followed by Scheffé’s post hoc test was performed. Otherwise, a nonparametric Mann–Whitney U test was employed, with the Bonferroni correction for multiple comparisons between samples applied. For comparisons between pairs of means (pairwise comparisons), Student’s *t*-tests were employed.

### Data availability

RNA-seq data were uploaded to the Gene Expression Omnibus under accession number GSE253964.

## Acknowledgments

This work is dedicated to the memory of mentor and friend, José Ramón Díaz-Ruiz Alba. The research was funded by grants PID2019-109304RB-I00 and PID2022-137691OB-I00 from the Spanish Ministry of Science and Innovation (MCIN/AEI/10.13039/501100011033), and by “ERDF A way of making Europe”. Khouloud Necira was funded by the Ministry of Higher Education and Scientific Research of Tunisia, grant numbers 2019-BALT-1077 and 2019-BALT-1079. We thank Dr. K. Kalantidis (University of Crete and IMBB/FoRTH,Greece) and Dr. J.J. López-Moya (Centre for Research in Agricultural Genomics, Spain) for providing *N. benthamiana* Dcli and Ago2KO transgenic lines, respectively.

## LEGENDS

Supplementary Table 1. Differentially expressed *N. benthamiana* genes performed for paired-comparisons made between treatments.

Supplementary Table 2. KEGG analysis of differentially expressed *N. benthamiana* genes performed for paired-comparisons made between treatments.

Supplementary Table 3. Differentially expressed *N. benthamiana* genes classified in MAPK signaling, Plant-pathogen interaction and Plant hormone signal transduction in the different comparisons.

